# Extreme mito-nuclear discordance within Anthozoa, with notes on unique properties of their mitochondrial genomes

**DOI:** 10.1101/2022.10.18.512751

**Authors:** Andrea M. Quattrini, Karen Snyder, Risa Purow-Ruderman, Isabela G.L. Seiblitz, Johnson Hoang, Natasha Floerke, Nina I. Ramos, Herman H. Wirshing, Estefanía Rodriguez, Catherine S. McFadden

## Abstract

Whole mitochondrial genomes are often used in phylogenetic reconstruction. However, discordant patterns in species relationships between mitochondrial and nuclear phylogenies are commonly observed. Within Anthozoa (Phylum Cnidaria), mitochondrial-nuclear discordance has not yet been examined using a large and comparable dataset. Here, we used data obtained from target-capture enrichment sequencing to assemble and annotate mitochondrial genomes and reconstruct phylogenies for comparisons to phylogenies inferred from 100s of nuclear loci obtained from the same samples. The datasets comprised 108 hexacorals and 94 octocorals representing all orders and >50% of extant families. Results indicated rampant discordance between datasets at every taxonomic level. This discordance is not attributable to substitution saturation, but rather likely caused by recent and ancient introgressive hybridization and selection. We also found strong purifying selection across the mitochondrial genomes, cautioning their use in analyses that rely on assumptions of neutrality. Furthermore, unique properties of the mitochondrial genomes were noted, including genome rearrangements and the presence of *nad5* introns. Specifically, we note the presence of the homing endonuclease in ceriantharians. This large dataset of mitochondrial genomes further demonstrates the utility of off-target reads generated from target-capture data for mitochondrial genome assembly and adds to the growing knowledge of anthozoan evolution.

## Introduction

Mitochondrial (mt) genes have a long history of use for phylogenetic reconstruction in animals [1], and the relative ease with which complete mt genomes can now be obtained has fueled an increase in their use to resolve phylogenetic relationships within many groups [2-4]. Animal mt genomes typically include a highly conserved set of protein-coding genes with few non-coding intergenic regions; are inherited uniparentally without undergoing recombination; and in many cases have rates of substitution that may be an order of magnitude higher than those of the nuclear genome [5]. While these properties are advantageous for phylogenetic reconstruction, they may also generate phylogenetic signals that differ from those of the nuclear genome. Discordance between nuclear and mt gene phylogenies is common and can result from biological processes such as introgression or incomplete lineage sorting (ILS) that act differently on mt vs. nuclear genomes [e.g., 6-8]. Alternatively, apparent mito-nuclear discordance can arise from inaccurate estimation of phylogenies due to low statistical power, poor model fit or taxon sampling issues [8]. Recent advances in computational models and increased taxon sampling of both mt and nuclear genomes have allowed these alternative sources of discordance to be evaluated in several well-sampled vertebrate taxa [6,8]. Studies have concluded that mito-nuclear discordance more often arises from biological processes such as introgression and ILS and persists even when factors that lead to inaccurate phylogenetic estimation have been addressed [6-8].

Phylogenies of anthozoan cnidarians (e.g., corals and sea anemones) reconstructed from mt genes or genomes have often recovered relationships within and among orders that differ from those inferred from both nuclear genes and morphology. The mt genomes of these non-bilaterian metazoans have several unusual properties that are not found in bilaterians [9] that may contribute to mito-nuclear discordance in this group. For example, the mt genomes of class Hexacorallia (e.g., sea anemones, scleractinian corals and black corals) encode the standard 13 protein-coding genes found in bilaterians, but only two tRNAs (trnW, trnM) [10-14]. Many hexacorals have group I introns in *nad5* or *cox1* [10-13], and the latter gene may have a LAGLI-DADG type homing endonuclease encoded within it [13]. The ceriantharian tube anemones have multipartite linear mt genomes [15]. All members of class Octocorallia (e.g., soft corals, gorgonians and sea pens) have just a single tRNA (trnM), but with only one known exception (i.e., a member of genus *Pseudoanthomastus*; [16]) their mt genomes include an additional protein-coding gene that encodes the DNA mismatch repair protein, *mtMutS* [17]. At least one sea pen has a bipartite circular mt genome [18], and other octocoral lineages have undergone frequent rearrangements (inversions) of gene order by a mechanism that appears to involve intramolecular recombination [19-21].

The unusual property of anthozoan mt genomes that has most impacted their utility for phylogenetic reconstruction is, however, the rate at which they evolve. Unlike bilaterian mt genomes that tend to evolve 5-10X faster than the nuclear genome [22-23], anthozoan mt genes typically evolve 10-100X slower than nuclear genes [24]. As a result, mt genes that have been widely used in bilaterians for barcoding, species-level phylogenetic analyses and phylogeography are often invariant within—and sometimes between—anthozoan genera [25-26]. These slow rates of mt gene evolution have, however, increased the potential utility of mt genes for reconstructing deep phylogenetic relationships among the families and orders of Anthozoa, a group of organisms that last shared a common ancestor in the pre-Cambrian [27-28]. Nonetheless, phylogenies of Anthozoa reconstructed from complete mt genomes (or their protein-coding genes) have often been incongruent with other sources of morphological and phylogenomic evidence. The most notable of these discrepancies has been a lack of support for the monophyly of the anthozoan classes, Hexacorallia and Octocorallia. Mitochondrial phylogenies have often placed Octocorallia sister to the cnidarian sub-phylum Medusozoa [4, 21, 29, 30], despite the very strong morphological and life-history evidence for the monophyly of Anthozoa [see 31], which has also been confirmed in several phylogenomic studies [32-33]. Moreover, in some of these same analyses Hexacorallia has been recovered outside of Cnidaria, as the sister to a clade of sponges [4, 34]. Mitochondrial gene phylogenies have also recovered Ceriantharia (tube anemones) sister to the rest of Anthozoa [15, 30, 35] rather than within Hexacorallia as supported by genomic-scale studies [27-28, 32]. In addition, previous studies have suggested that Scleractinia is paraphyletic with Corallimorpharia [4, 12, 36]) and have differed from nuclear gene phylogenies in the placement of the orders Actiniaria, Zoantharia and Antipatharia and in the relationships among the major clades of Scleractinia [37-38]. Within Octocorallia, mt genes and/or genomes have provided little statistical support for the deepest nodes in either of the two major clades that have been recognized [29, 30, 39, 40].

Explanations that have been proposed to explain the incongruence between mt and nuclear or morphological phylogenies of Anthozoa include substitution saturation of the mt genome [21, 36, 41], rate heterogeneity between the major lineages [29], and long branch attraction (LBA) due to the combined effects of rate heterogeneity and incomplete or biased taxon sampling [34]. Most mt genome phylogenies and phylogenomic analyses of anthozoans published to date have been taxon-sparse, often omitting entire orders [29, 32, 33] or have drawn comparisons between topologies generated from completely different taxon sets [41]. As a result, it is still unclear if the source of incongruence between mt and nuclear gene phylogenies of anthozoans is simply an artifact of incomplete, biased and incomparable taxon sampling or if the evolutionary signal present in anthozoan mt genomes does indeed differ from that of the nuclear genome.

Recent advances in phylogenomic methods and technologies have facilitated the ability to obtain complete mt genomes while simultaneously generating sequence reads for thousands of nuclear genes. In particular, target-enrichment methods used to sequence ultraconserved elements (UCEs) and exonic regions of the nuclear genome can recover complete or near-complete mt genomes as off-target reads [3]. Comparisons of mt vs nuclear gene phylogenies from the same set of taxa (often the same individuals) facilitate investigation of the causes of mito-nuclear incongruence by eliminating artifacts that may be caused by unequal or different taxon sampling.

In recent phylogenomic analyses of Anthozoa based on UCEs and exons [27-28], complete or near-complete mt genomes were recovered for a majority of the taxa sequenced. Here we used the complete set of mt protein-coding sequences to reconstruct the phylogenies of the Octocorallia and Hexacorallia classes and compared those to nuclear gene phylogenies generated for the same set of individuals. The dataset comprised a total of 202 species representing all orders and >50% of extant families. With this comparable dataset, the impacts of sampling biases were removed and we were able to robustly explore whether incongruence is related to evolutionary signal. New findings on the unique properties of the recovered mt genomes are also noted.

## Methods

### Target-Enrichment Analyses

UCE and exon loci were target enriched and bioinformatically extracted from high-throughput sequencing data as described in Quattrini et al. [42] and [27] using the anthozoa-v1 baitset [42]. Briefly, raw reads were cleaned using illumiprocessor [43] and Trimmomatic v 0.35 [44] and then assembled using either Spades v 3.1 ([45]; with the --careful and --cov-cutoff 2 parameters) or Trinity v. 2.0 [46]. The phyluce pipeline was then used as described in the online tutorials (https://phyluce.readthedocs.io/en/latest/tutorials/tutorial-1.html) with some modifications (see supplemental code in 27, 42). Using phyluce, 75% and 50% taxon-occupancy matrices were created for each nuclear locus, aligned with MAFFT v7.130b [47], and loci were concatenated (*phyluce_align_format_nexus_files_for_raxml*) separately for hexacorals (n=108) and octocorals (n=94).

### Mitochondrial Genome Analyses

Whole and partial mt genomes were extracted from the off-target reads in the target-enrichment sequencing data. Mitochondrial genomes were extracted and assembled in three ways. First, we used blastn to find whole or partial genomes in the Trinity or Spades assemblies and then extracted those as fasta sequences. Second, we used Novoplasty v 2.6 [48] to assemble mt genomes using the adapter-trimmed paired-end reads. Seed files were used to help assemble each species and consisted of *cox1* sequences downloaded from GenBank for the species of interest or a closely-related species. Third, Geneious Prime 2020 (https://www.geneious.com) was used for genomes that were difficult to assemble with Spades and Novoplasty. Individual mt loci from closely related taxa, either *mtMutS, cox1* or 16S, were used as seeds to initiate and guide assemblies.

Following mt genome assembly, fasta files were uploaded to Mitos2 ([49], http://mitos2.bioinf.uni-leipzig.de) for annotation (translation code=4). For further analyses, we used only species whose mt genomes were represented by at least 50% of the protein coding genes (hexacorals n=108, octocorals n=94, Suppl. Table 1), except that we included five ceriantharians with low mitogenome recovery (e.g., for 15-53% of genes recovered for each species). Protein-coding genes were then each aligned separately using MAFFT v7.130b [47] and adjusted by eye to ensure the sequences were in frame. Loci were then concatenated with *phyluce_align_concatenate_alignments*. Mitochondrial genomes of hexacorals were deposited in GenBank under BioProject #XXX.

Some mt genomes for which we had corresponding nuclear data could not be easily assembled, or were published in previous studies, and so sequences were downloaded from GenBank and subsequently used in our analyses (Suppl. Table 1). We used mt data from GenBank for 26 hexacorals; 16 of these were of the same individuals used in our study. All octocoral mt genomes were also assembled concurrently in another study [16] and added to GenBank by those authors.

### Phylogenomic Analyses

Removing loci that are saturated can improve phylogenomic analyses [50]. Therefore, we ran saturation tests on each of the different locus datasets using Phylomad [51]. For nuclear loci, we ran saturation tests using models of entropy on all sites and only on those that had no missing data in each locus alignment. Datasets are denoted hereafter as LR (low risk loci) and LRM (low risk loci with no missing data in saturation test). For the mt data, we ran saturation tests on sites with no missing data for the concatenated alignment. Loci with substitution saturation were removed and then various datasets were used for further phylogenetic analyses (Suppl.Table 2, Table 1).

**Table 1.** Summary statistics for different alignment datasets.

| Dataset | Taxon<br>Occupancy | #<br>Loci | Alignment<br>Size | # BS<br>>95 | # BS<br>95-80 | # BS<br>< 80 | # Sh<br>>95 | # Sh<br>95-80 | # Sh<br><80 |
| --- | --- | --- | --- | --- | --- | --- | --- | --- | --- |
| **Hexacorallia** |  |  |  |  |  |  |  |  |  |
| ASTRAL | 75% | 222 | NA | 95* | 2* | 9* | NA | NA | NA |
| ASTRAL LR | 75% | 149 | NA | 89* | 4* | 13* | NA | NA | NA |
| ASTRAL LRM | 75% | 95 | NA | 92* | 4* | 10* | NA | NA | NA |
| 75 | 75% | 222 | 85327 | 100 | 3 | 3 | 101 | 1 | 5 |
| 75LR | 75% | 149 | 66567 | 100 | 3 | 3 | 100 | 1 | 4 |
| 75LRM | 75% | 95 | 38534 | 93 | 7 | 6 | 96 | 2 | 8 |
| 50 | 50% | 756 | 246027 | 103 | 2 | 1 | 103 | 2 | 1 |
| 50LR | 50% | 545 | 196468 | 103 | 1 | 2 | 103 | 1 | 2 |
| 50LRM | 50% | 380 | 126183 | 99 | 6 | 1 | 100 | 3 | 3 |
| Mito | 94-98% | 13 | 12465 | 94 | 7 | 5 | 83 | 11 | 12 |
| **Octocorallia** |  |  |  |  |  |  |  |  |  |
| ASTRAL | 75% | 621 | NA | 83* | 4* | 4* | NA | NA | NA |
| ASTRAL LR | 75% | 575 | NA | 83* | 1* | 7* | NA | NA | NA |
| ASTRAL LRM | 75% | 408 | NA | 76 | 7 | 8 | NA | NA | NA |
| 75 | 75% | 621 | 305233 | 82 | 3 | 6 | 88 | 2 | 1 |
| 75LR | 75% | 575 | 290130 | 85 | 2 | 4 | 87 | 2 | 2 |
| 75LRM | 75% | 408 | 213477 | 80 | 8 | 4 | 85 | 4 | 2 |
| 50 | 50% | 1252 | 555701 | 82 | 5 | 4 | 87 | 2 | 2 |
| 50LR | 50% | 1161 | 529092 | 85 | 2 | 4 | 85 | 1 | 5 |
| 50LRM | 50% | 827 | 385627 | 85 | 4 | 2 | 86 | 3 | 2 |
| Mito | 77-100% | 14 | 16176 | 69 | 14 | 8 | 69 | 12 | 10 |

BS=bootstrap, SH= Shimodaira-Hasegawa approximate likelihood ratio test, LRM=low risk loci only with no missing data in saturation test, LR=low risk loci only while allowing for missing data in saturation test, mito=mitochondrial alignment, \*=posterior probability

Selection tests were conducted using codon-based models in codeml within PAML v. 4 [52]. The one ratio model (M0) was run on the mt alignment only for both octocorals and hexacorals. This allowed us to estimate average omega (*dN/dS*) and kappa (*ts/tv*) values across all branches in the corresponding mt phylogenies. Omega values =1 indicate the locus is evolving neutrally, values >1 indicate positive selection and values <1 indicate negative or purifying selection. Higher kappa values indicate transition relative to transversion bias.

Phylogenomic analyses were conducted using maximum likelihood in IQTree v 2.1 [53] on each of the concatenated datasets (Table 1). We ran partitioned analyses on the different datasets using the best model for each locus chosen with ModelFinder [54]. Ultrafast bootstrapping (-bb 1000, [55]) and the Sh-like approximate likelihood ratio test (-alrt 1000, [56]) were conducted as well as site-concordance factors [57]. A species tree analysis was also conducted using ASTRAL III v 5.7, which is statistically consistent under a multispecies coalescent model [58]. Gene trees were constructed in IQTree using the best fit model of evolution selected with ModelFinder for each gene. We used the 75% taxon-occupancy data matrices for each class. Treeshrink [59] was used to remove long branches, and the newick utility, nw_ed, was used to remove branches with <30 % bootstrap support prior to running IQTree. Site concordance factors were also calculated on the species tree using the concatenated alignment. The phylogenetic relationship of *Renilla muelleri* to other octocorals was spuriously placed in some phylogenies. Because this species is well-supported in Pennatuloidea, we pruned this species from all phylogenetic trees using the R *phytools* package [60].

Following phylogenetic inference, we conducted Robinson-Foulds distance (R-F, [61]) tests using IQTree v2.1. R-F distances were calculated between all pairs of hexacoral unrooted trees and all pairs of octocoral unrooted trees. The two most congruent mt and nuclear trees based on maximum likelihood were determined based on the smallest R-F distances for both hexacorals and octocorals and plotted. In cases where R-F distances were the same, we chose the topology with the most bs support values > 95%. Hexacorals were rooted at the Ceriantharia based on prior phylogenomic studies of the phylum Cnidaria [32-33] and Scleralcyonacea was rooted to Malacalcyonacea based on prior phylogenomic studies [27-28, 40]. All code can be found in Suppl. File S1 and all trees and alignments can be found on figshare.

## Results

### Mitogenome Assemblies

Herein we assembled complete or near complete mt genomes of 75 hexacorals from the following orders: Actiniaria, Antipatharia, Ceriantharia, Corallimorpharia, and Scleractinia. Ceriantharian mt genomes were difficult to assemble. Out of five ceriantharians, none had complete mt genomes and only two genes were found for one species (*Ceriantheomorphe brasiliensis)*. Only one species, *Botruanthus mexicanus*, had a near complete genome assembly. We confirmed the presence of Group I introns (Suppl. Table 3) in many taxa. In Actiniaria, Antipatharia, Zoantharia, and *Relicanthus daphneae*, two protein coding genes, *nad1* and *nad3*, were found inserted as introns within *nad5*. Ten protein coding genes were found in the *nad5* intron of most scleractinians, with the exception of *Caryophyllia arnoldi* in which we found only seven protein-coding genes and *rns* within the *nad5* intron. In Corallimorpharia, 10 protein-coding genes were in the *nad5* intron of *Corallimorphus profundus*, and for the rest of the corallimorpharians (*Rhodactis osculifera, Discosoma carlgreni*, and *Ricordea florida)*, all genes but *trnW* were in the *nad5* intron. Another group 1 intron that encodes a homing endonuclease from the LAGLI-DADG family was present in *cox1* of some hexacorals. We confirmed the presence of this endonuclease in 24% of actiniarians, 28% of scleractinians, 17% of antipatharians, and 100% of corallimorpharians (Suppl. Table 3). We also documented this intron in two species of Ceriantharia, *Botruanthus mexicanus* and *Ceriantheomorphe brasiliensis*.

Of the complete (or near complete) mt genomes of hexacorals assembled in this study, only three species displayed gene order rearrangements relative to other taxa in their respective orders (Suppl. Table 3). Within Actiniaria, only one species sequenced, *Alicia sansibarensis*, exhibited a mt genome rearrangement with *cox2-nad4-nad6-cob* inserted prior to *atp8* instead of between *nad6* and *rns*. Of the scleractinians, *Caryophyllia arnoldi* had a genome rearrangement with the *cob-nad2-nad6* gene block inserted after the 3’ end of *nad5* instead of within the *nad5* intron. The mt genome of *Madrepora oculata* also had a gene rearrangement, with a switch in the order of *cox2* and *cox3* compared to all other scleractinians. *Corallimorphus profundus* also had a different genome rearrangement compared to *R. osculifera. D. carlgreni*, and *R. florida* (Suppl. Fig. 3). *Corallimorphus profundus* had 10 protein-coding genes and *rns* within the *nad5* intron. In contrast, *R. osculifera, D. carlgreni* and *R. florida* have all other genes but *trnW* within the *nad5* intron.

### Alignment Summary

For hexacorals, concatenated nuclear locus alignments across 50-75% taxon-occupancy datasets ranged from 38,534 to 246,027 bp with 95 to 756 loci in each dataset (Table 1). For each hexacoral species, locus recovery ranged from 303 to 1,156, with overall few loci (342 to 589) recovered in ceriantharians (Suppl Table 1). For octocorals, concatenated nuclear locus alignments across 50-75% taxon-occupancy datasets ranged from 213,477 to 555,701 bp with 408 to 1,252 loci (Table 1). For each octocoral species, 604 to 1,275 loci were recovered (Suppl. Table 1).

All 13 protein-coding genes were included in the alignment for 79% of all hexacoral species (Suppl. Tables 1 and 3). The hexacoral mt genome alignment containing the 13 protein-coding genes was 12,465 bp, and for each gene at least 94-98% of the species were represented. For octocorals, all 14 protein-coding genes were included in the alignment for 80% of species. The octocoral mt genome alignment was 16,176 bp, and for each gene 96-100% of the species were represented, except for *mtMutS. mtMutS* was included for only 77% of the species as for some speices it was highly incomplete or highly divergent from the other species.

Selection tests on the mt genome alignments indicated that the mt genomes are under strong purifying selection. The omega value (*dN/dS*) for hexacorals was 0.10 while the value for octocorals was 0.14. The kappa value (ts/tv) for hexacorals was 2.7, whereas in octocorals it was higher at 3.9. Saturation tests conducted using PhyloMad indicated that neither the hexacoral nor octocoral mt alignment was under saturation as indicated by entropy tests (Suppl. Fig. 1, Suppl. Table 2). For nuclear-locus datasets, 8-50% of the loci in each dataset had a high risk of substitution saturation. Hexacorals tended to have more saturated loci, with 30-50% saturated loci per dataset whereas octocorals had a lower number, with 8-35% of saturated loci per dataset.

### Mito-nuclear Discordance

#### Hexacorallia

Overall, all phylogenies constructed for Hexacorallia were well supported (Table 1). Among all nuclear trees constructed with ASTRAL and IQTree, 83 to 97% of nodes (106 total nodes) on each tree had higher than 95% ultrafast bootstrap (bs) values, posterior probabilities (pp), and SH-aLRT values. Similarly, the mt genome tree was well supported with 78-89% of nodes having higher than 95% ultrafast bs values and SH-aLRT values.

There were some differences among all hexacoral phylogenies, but nuclear phylogenies constructed with ASTRAL and IQTree were mostly congruent with one another (pairwise R-F distances=4-28). The R-F distances between the hexacoral mt genome tree and the nuclear trees, however, were much larger, ranging from 56 to 68. The mt genome tree was most similar to the ASTRAL species trees (pairwise R-F distances=56-60) compared to the maximum likelihood phylogenies (pairwise R-F distances=62-66); there were three maximum likelihood phylogenies that were all equally congruent with the mt genome phylogeny (RF distances=62). The tree with 50% data occupancy and highly saturated loci removed (50LR) while allowing for missing data in the saturation test had the highest BS support values. There were a few species on long branches in the mt genome tree, but not the nuclear tree, including the zoantharians *Nanozoanthus harenaceus* and *Microzoanthus occultus* and the scleractinian *Paraconotrochus antarcticus* (see Suppl. Files).

Although there were several differences among shallow nodes in all topologies, two major differences were apparent at deep nodes (Fig. 1). First, the relationship of Zoantharia and Actiniaria to other orders differed among topologies. In the mt genome tree, Actiniaria was sister to all other hexacoral orders except Ceriantharia (bs=100, SHaLRT=100, sCF=82). This same relationship was also recovered in the ASTRAL species trees (bs=100, SHaLRT=100, sCF=54-56) and the 75LRM tree (bs=100, SHaLRT=100, sCF=56, Fig. 1, see Suppl. Files) analyses of the nuclear dataset. In contrast, in the majority of maximum likelihood trees for the nuclear dataset, Zoantharia diverged earlier than Actiniaria (bs=100, SHaLRT=100, pp=100, sCF=52-84). Second, the relationship of *Relicanthus daphneae* to other orders differed among phylogenies. In the mt genome phylogeny, *R. daphneae* was sister to the zoantharians (UF=88, SHaLRT=77, sCF=42). In the majority of nuclear phylogenies, *R. daphneae* was recovered with variable support (bs >84, SHaLRT >75, pp>41, sCF >32), as sister to Antipatharia-Corallimorpharia-Scleractinia.

**Figure 1.**
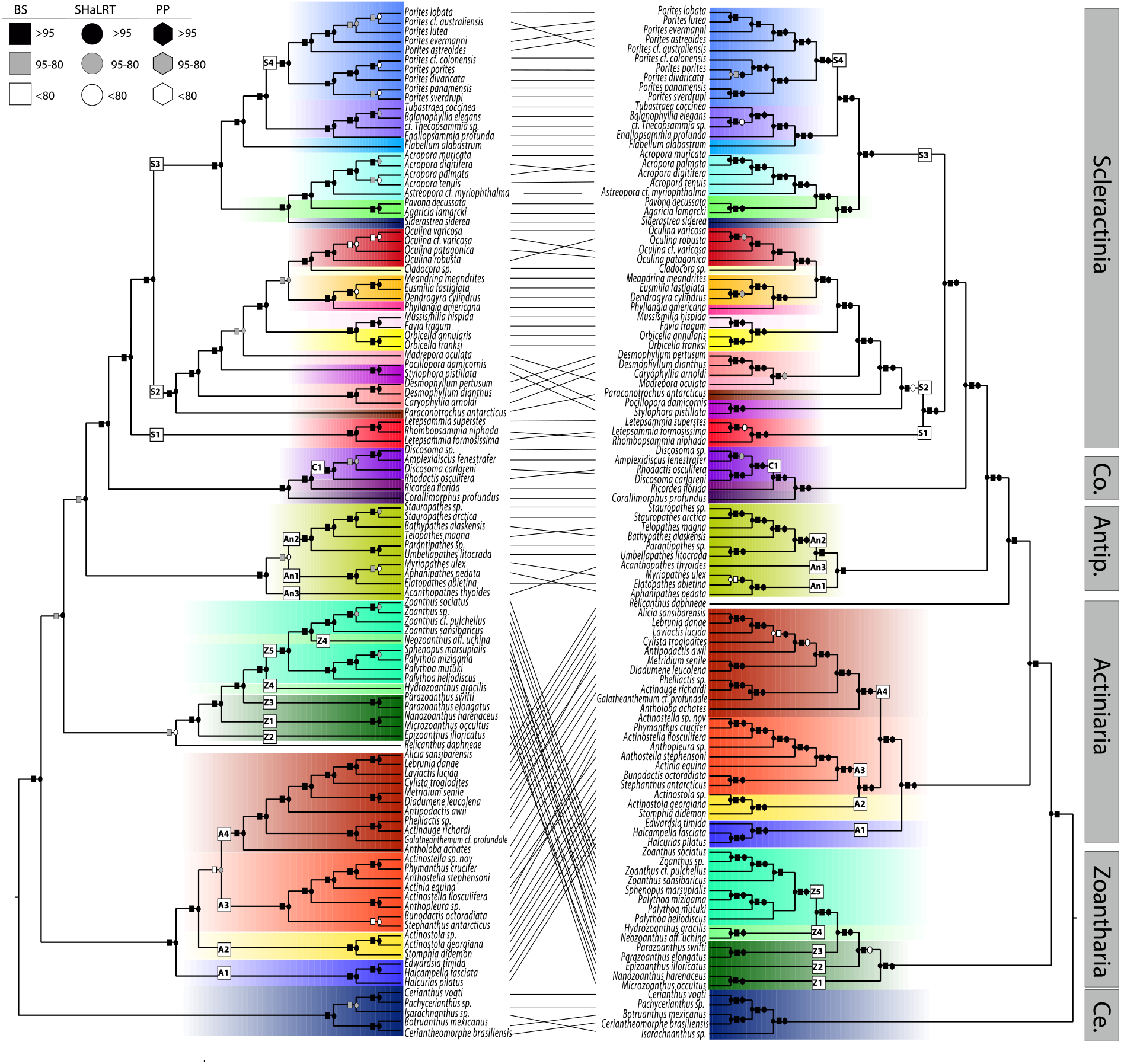
Maximum likelihood tree of Hexacorallia inferred from (left) mitochondrial and (right) nuclear loci (50% taxon occupancy, no loci with substitution saturation as denoted in saturation tests with missing data included). Numbered squares on branches identify clades discussed in Results.

There were also some differences between mt genome and nuclear phylogenies within each hexacoral order. Within Scleractinia, there were differences among trees at the shallow nodes, including relationships among species of *Porites* (S4 clade), but the major difference was the placement of the family Micrabaciidae (S1). Nuclear phylogenies all strongly support that this family is sister to the Robust/Vacatina clade (S2) of Scleractinia (bs=100, SHaLRT=100, pp=97-100, sCF=37-40). In contrast, in the mt genome phylogeny Micrabaciidae (S1) was recovered as sister to all other Scleractinia (S2+S3) with strong support (bs=100, SHaLRT=99, sCF=33). Within Actiniaria, the position of the superfamily Actinostoloidea (A2) differed between mt genome and nuclear phylogenies. This superfamily was sister to the superfamilies Metridioidea+Actinioidea (A4+A3) in the mt genome phylogeny (bs=100, SHaLRT=100, sCF=48.6) whereas it was sister to the superfamily Actinioidea (A3) in all nuclear phylogenies (bs=100, SHaLRT=99-100, pp=100, sCF=37-38). Within Antipatharia, the position of *Acanthopathes thyoides* (A3) differed between mt genome and nuclear phylogenies. This species was sister to all other antipatharians (An1+An2) in the mt genome phylogeny (bs=100, SHaLRT=100, sCF=72) whereas it was sister to the family Schizopathidae (An2) in the majority of nuclear phylogenies (bs=76-100, SHaLRT=23-100, pp=91-98, sCF=34-37), except for the 50LRM and ASTRAL LRM topologies (bs=100, SHaLRT=100, pp=100, sCF=65-67), which matched the mt genome tree. Within Zoantharia, the placement of *Epizoanthus illoricatus* (Z2) and *Neozoanthus* aff. *uchina* (Z4 in part) differed among mt genome and nuclear phylogenies. In the mt genome tree, *E. illoricatus* (Z2) was sister to the rest of the zoantharians (bs=100, SHaLRT=100, sCF=66), whereas in all nuclear phylogenies, *Nanozoanthus harenaceus* and *Microzoanthus occultus* (Z1) were sister to the rest of the zoantharians (bs=100, SHaLRT=100, pp=100, sCF=37-38). *Neozoanthus*. aff. *uchina* (Z4 in part) was sister to the family Zoanthidae (Z5) in the mt genome phylogeny (bs=100 SHaLRT=100, sCF=49) whereas it was sister to *Hydrozoanthus gracilis* (Z4) in all nuclear phylogenies (bs=100 SHaLRT=100, pp=100, sCF=48-51). Within Corallimorpharia, there were differences within the Discosomidae family (C1) with *R. osculifera* sister to *D. carlgreni* in the nuclear phylogeny yet sister to the remaining discosomids in the mt genome phylogeny.

#### Octocorallia

Nuclear gene phylogenies for Octocorallia were in general well supported. Among all nuclear trees constructed with ASTRAL and IQTree, 83 to 96% of nodes (91 total nodes) on each tree had higher than 95% ultrafast bootstrap (bs) values, posterior probabilities (pp), and SH-aLRT values. In contrast, mt genome trees for Octocorallia were not as well supported with only 76% of nodes having higher than 95% ultrafast bs and SHaLRT values.

Nuclear phylogenies constructed with ASTRAL and IQTree were somewhat congruent with one another (pairwise R-F distances=4-36, Table 2, see Suppl. Files). The R-F distances between the octocoral mt genome tree and the nuclear trees, however, were much larger, ranging from 60 to 72. Octocoral mt genome trees were somewhat more similar to the maximum likelihood phylogenies (pairwise R-F distances=60-68) as compared to the ASTRAL trees (pairwise R-F distances=68-72).The most similar tree to the mt genome phylogeny was constructed with a 75% taxon occupancy data matrix with highly saturated loci removed and no missing data in the saturation test (75LRM). In general, branch lengths were much different between mt genome and nuclear trees. In the mt genome tree, seven species were on very long branches (*Muricella* sp., *Leptophyton benayahu, Tenerodus fallax, Cornularia pabloi, Pseudoanthomastus* sp., *Erythropodium caribaeorum*, and *Melithaea erythraea*), a pattern not recovered in nuclear phylogenies (see Suppl. Files).

**Table 2.** Pairwise Robinson-Foulds distances between hexacoral topologies and octocoral topologies.

|  | ASTRAL | ASTRAL<br>LR | ASTRAL<br>LRM | 75 | 75LR | 75LRM | 50 | 50LR | 50LRM | Mito |
| --- | --- | --- | --- | --- | --- | --- | --- | --- | --- | --- |
| **Hexacorallia** |  |  |  |  |  |  |  |  |  |  |
| ASTRAL | 0 |  |  |  |  |  |  |  |  |  |
| ASTRAL LR | 4 | 0 |  |  |  |  |  |  |  |  |
| ASTRAL LRM | 12 | 12 | 0 |  |  |  |  |  |  |  |
| 75 | 18 | 20 | 28 | 0 |  |  |  |  |  |  |
| 75LR | 16 | 16 | 26 | 6 | 0 |  |  |  |  |  |
| 75LRM | 26 | 24 | 26 | 20 | 20 | 0 |  |  |  |  |
| 50 | 14 | 14 | 20 | 14 | 14 | 22 | 0 |  |  |  |
| 50LR | 16 | 14 | 24 | 8 | 4 | 18 | 12 | 0 |  |  |
| 50LRM | 26 | 24 | 24 | 18 | 18 | 22 | 12 | 14 | 0 |  |
| Mito | 58 | 60 | 56 | 62 | 62 | 66 | 64 | 64 | 62 | 0 |
| **Octocorallia** |  |  |  |  |  |  |  |  |  |  |
| ASTRAL | 0 |  |  |  |  |  |  |  |  |  |
| ASTRAL LR | 6 | 0 |  |  |  |  |  |  |  |  |
| ASTRAL LRM | 14 | 12 | 0 |  |  |  |  |  |  |  |
| 75 | 28 | 32 | 30 | 0 |  |  |  |  |  |  |
| 75LR | 34 | 36 | 36 | 6 | 0 |  |  |  |  |  |
| 75LRM | 30 | 30 | 28 | 14 | 20 | 0 |  |  |  |  |
| 50 | 24 | 28 | 30 | 10 | 16 | 24 | 0 |  |  |  |
| 50LR | 28 | 32 | 32 | 4 | 10 | 18 | 6 | 0 |  |  |
| 50LRM | 28 | 32 | 32 | 4 | 10 | 18 | 10 | 8 | 0 |  |
| Mito | 72 | 70 | 68 | 68 | 66 | 60 | 68 | 66 | 68 | 0 |

LRM=low risk loci only with no missing data in saturation test, LR=low risk loci only with missing data in saturation test, mito=mitochondrial alignment, 75=75% taxon occupancy dataset, 50=50% taxon occupancy dataset

Numerous differences were apparent among the octocoral mt genome and nuclear phylogenies (Fig. 2). Within the order Scleralcyonacea, the placement of Pennatuloidea+Ellisellidae (clade S1) differed. In the mt genome tree this clade was sister to the Keratoisididae+Primnoidae+Chrysogorgiidae (S2) and Helioporidae (S3) clades (bs=90, SHaLRT=100, sCF=29). In two maximum likelihood trees (50, 50LRM) and all ASTRAL trees (bs=100, SHaLRT=100, pp=93-100, sCF=36), it was sister either to clades S3+S4 (bs=53-100, SHaLRT=25-99, sCF=33-36) or to clades S2+S3+S4. *Cornularia pabloi* also changed positions, diverging later (sister to clade S3) in the mt genome phylogeny (bs=90, SHaLRT=95, sCF=28) as compared to all nuclear phylogenies where it was placed sister to all other scleralcyonaceans (bs=100, SHaLRT=100, pp=100, sCF=37). *Parasphaerasclera valdiviae* was an early-diverging lineage and sister to all other scleralcyonaceans in the mt genome phylogeny (bs=100, SHaLRT=100, sCF=63) whereas it was sister to family Coralliidae in the nuclear phylogeny (bs=100, SHaLRT=100, pp=100, sCF=37). Helioporidae (S3) was recovered as sister to clade S4 in the maximum likelihood nuclear phylogenies (bs=99-100, SHaLRT=99, sCF=36) but sister to clade S2 in the mt genome phylogeny (bs=90, SHaLRT=97, sCF=34) and the ASTRAL phylogenies, although the relationships in the species trees were poorly to moderately supported (pp=0.5-86, sCF=34). Family Keratoisididae was recovered as sister to Primnoidae in the mt genome phylogeny (bs=94, SHaLRT=96, sCF=32) and in one nuclear phylogeny (50LRM) but with poor support (bs=79, SHaLRT=47, sCF=32). In all other nuclear phylogenies, Keratoisididae was recovered sister to Chrysogorgiidae (bs=100, SHaLRT=100, pp=90-100, sCF=36-37).

**Figure 2.**
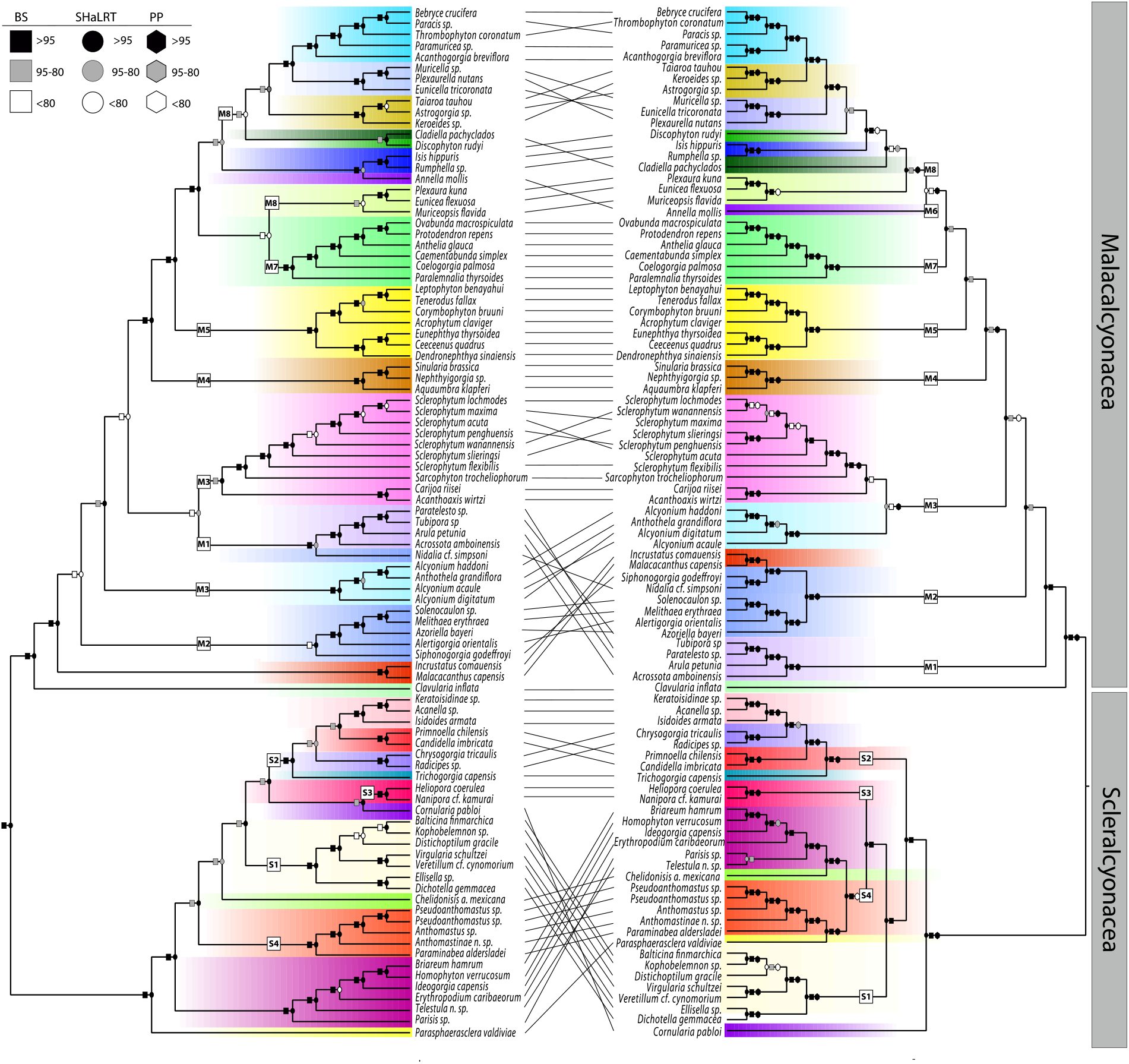
Maximum likelihood tree of Octocorallia inferred from (left) mitochondrial and (right) nuclear loci (75% taxon occupancy, no loci with substitution saturation as denoted in saturation tests without missing data). Numbered squares on branches identify clades discussed in Results.

Within Malacalcyonacea, several differences among phylogenetic relationships were noted, including some relationships among congeneric species. The Incrustatidae+Malacacanthidae clade was an early-diverging lineage and sister to most malacalcyonacean families (except for *Clavularia inflata)* in the mt genome tree (bs=100, SHaLRT=100, sCF=50), but these families diverged later as part of the M2 clade in the nuclear phylogenies. The Tubiporidae+Arulidae clade (M1) was sister to all malacalcyonaceans (except for *C. inflata)* in the nuclear phylogeny (75LRM). In the mt genome phylogeny, it included *Nidalia* and was sister to the Sarcophytidae+Carijoidae (clade M3a, bs=72, SHaLRT=86, sCF=32). An Anthogorgiidae+Eunicellidae+Plexaurellidae clade (M8a) was sister to Paramuriceidae (M8c) in the mt genome phylogeny (bs=99, SHaLRT=95, sCF=35). In contrast, the Keroeidae+Taiaroidae+Astrogorgiidae clade (M8b) was sister to the Paramuriceidae (M8c) in the nuclear phylogenies (bs=99-100, SHaLRT=99-100, pp=100, sCF=34-42). Within Sarcophytidae, relationships differed among species between mt genome and nuclear phylogenies.

## Discussion

### Mitochondrial Genome Properties

Utilizing a total of 202 complete or near-complete mitochondrial (mt) genomes, we were able to examine mito-nuclear discordance within the Anthozoa and explore the unique mt genome properties of all orders belonging to this sub-phylum of Cnidaria. In addition to the mt genomes newly assembled here, most of the previously published mt genomes [16, 38, 62] that we included in our analyses had been assembled from the raw sequence data from Quattrini et al. [27, 42]. This large dataset of mt genomes further demonstrates the utility of off-target reads generated from target-capture data for the assembly of mt genomes and adds to the growing knowledge of mt genome evolution within the sub-phylum Anthozoa.

Although group I introns have been previously recorded in hexacorals [10-14, 62-65], we note their pervasiveness across the group. A *nad5* intron of at least two protein-coding genes and up to all 13 is present in the majority of hexacoral families. From our data, it also appears that this intron is present in Ceriantharia, however, this needs further confirmation as we had difficulties assembling mt genomes in that order. The other group I intron that encodes a homing endonuclease from the LAGLI-DADG family is present in *cox1* in many hexacorals. Both gains and/or losses of this gene have been previously noted in the hexacoral orders, Scleractinia [13], orallimorpharia [64], Actiniaria [65], and Zoantharia [62]. This endonuclease appears to be more common in some orders (Zoantharia, Corallimorpharia) than others (Scleractinia). Based on annotation from Mitos2, we also documented this intron in two ceriantharians. To our knowledge, this intron has not yet been documented in the order Ceriantharia. Based on its distribution across the phylogeny, the homing endonuclease, likely a result of horizontal transmission [13], has been gained and lost within Hexacorallia for several hundred million years, with origins dating to 300-400 MYA [27]. To date, no introns have been recorded from the Octocorallia.

Mitochondrial genome rearrangements within Anthozoa have been a topic of interest for over two decades, as species in this sub-phylum exhibit several gene order changes. However, within hexacorals, genome rearrangements are seemingly rare. Of the 102 complete (or near complete) mt genomes of hexacorals examined in this study, only 7% displayed gene order rearrangements relative to the canonical gene order within their taxonomic order; many of which have been described in prior studies [e.g., 12, 65, 66]. In contrast to hexacorals, octocorals have undergone gene rearrangements more frequently across their phylogenetic history [18-21, 67]. Of the 92 complete to near complete octocoral mt genomes used in this study, 21% had gene rearrangements. Brockman and McFadden [20] suggested that octocoral gene rearrangements evolve via inversions of conserved gene blocks (or intramolecular recombination) whereas hexacoral gene rearrangements are likely caused by gene shuffling. Additionally, they hypothesized that the presence of the mt mis-match repair protein, *mtMutS* (unique to Octocorallia) might play a role in mediating these gene inversions. A recent review by Johansen and Emblem [68] suggested that the large *nad5* intron that is ubiquitous in hexacorals (but absent from octocorals) perhaps stabilizes mt genome organization in that class. With the increasing availability and decreasing costs of high-throughput sequencing combined with new analytical methods for assembling and annotating mt genomes (e.g., MitoFinder, [3]), many new discoveries likely await regarding the mt genome evolution of anthozoan cnidarians.

### Mito-nuclear Discordance

Advances in genomic approaches have also facilitated comparisons of the phylogenetic histories of nuclear and mt genomes. This has allowed us to explore the patterns and underlying causes of mito-nuclear discordance. In both Hexacorallia and Octocorallia, we found a high-degree of mito-nuclear discordance at every level (i.e., order to species) even when comparing the mt phylogeny to the most similar nuclear phylogeny. At deep nodes in the phylogenies, the most apparent differences in the hexacoral phylogenies included the positions of the anemone groups Actiniaria, Zoantharia, and *R. daphneae*. Within octocorals, the most apparent differences at deep nodes were relationships among clades within the order Scleralcyonacea and among the early-diverging lineages within Malacalcyonacea. Discordance at deep nodes complicates interpretations of ancestral state reconstructions through deep time. In addition, this level of discordance causes concern for using just one source of sequence data (i.e. nuclear or whole mt genomes) for phylogenetic reconstruction, but also highlights how different datasets used in compliment present a unique opportunity to better understand the cause of the discordance from an evolutionary perspective.

Substitution saturation of mt genomes has been suggested to be the cause of mito-nuclear discordance in anthozoans [21,41]. Using entropy tests on our extensive dataset of ∼100 genomes in each class, we did not find evidence for substitution saturation. The entropy-based t statistic tests saturation on phylogenetically informative sites, is suitable for assessing misleading tree topologies, and it has several advantages, including: 1) it is robust across a range of confounding factors, including rate variation across sites; and 2) the negative influence of slowly-evolving sites is removed in the measurement of overall base composition [50]. Thus, our results might differ from prior studies that used other methods, particularly if slowly-evolving sites were not taken into account. Alternatively, the different results could be driven by the number (2-3X less) and choice of taxa used in prior phylogenetic studies. In contrast to mt genomes, we found that ∼10 to 50% of UCE and exon nuclear loci were saturated, depending on dataset. A recent study examining substitution saturation of UCE and exon loci across a range of taxa (e.g., hymenopterans, fishes, and crustaceans), also found similar numbers of saturated loci [50]. We removed UCE and exon loci with substitution saturation from the dataset prior to phylogenetic analysis, yet even so, the nuclear and mt topologies were quite incongruent. Therefore, substitution saturation is not the primary cause of the observed discordance among nuclear and mt phylogenies.

Introgression is another biological process that can result in discordance among nuclear and mt phylogenies. Within Anthozoa, introgressive hybridization has been suggested to be an important mechanism in generating species diversity [71-77]. Because mt genomes are maternally inherited and non-recombining, species or groups of species that have undergone past hybridization might be expected to have mt genomes that are more similar than their nuclear genomes [e.g., 78-79]. Using D-statistics and ABBA BABA tests, Quattrini et al. [75] determined that hybridization is an important mechanism in shaping diversity within the soft octocoral genus *Sclerophytum (*=*Sinularia*). Similarly, hybridization has been noted within multiple species in the scleractinian genus *Porites* [76-77]. Indeed, we found strong incongruence between mt and nuclear phylogenies within both genera. In combination, these results suggest that introgression might explain some of the incongruence, at least at the tips of the trees. Mitochondrial introgression is more likely and happens at a faster rate than nuclear introgression, cautioning the use of mt gene trees as accurate depictions of a species trees [79]. Future studies should consider explicitly testing for mt introgression in pairs or groups of taxa using, for example, ABBA-BABA tests and isolation with migration models [e.g., 80].

Studies that have tested for introgressive hybridization as a cause of discordance among mt and nuclear phylogenies have often focused on closely related species groups that have diverged in the recent past [e.g., 6, 8, 81-82]. Whether or not hybridization is the cause of incongruent relationships at nodes deeper in a phylogeny is more difficult to discern. However, ancient introgression of ghost lineages (e.g., extinct, unknown or unsampled lineages that remain in extant species likely due to ancient hybridization; [83]) could play a role in generating incongruence. Li and Wu [84] hypothesized that species with widespread distributions are likely to contain genetic components of ghost lineages. Marine invertebrates, such as anthozoans, fit that category. The case of the enigmatic giant deep-sea anemone *Relicanthus* particularly fits this scenario, so far being the only representative of its kind within hexacorals. We urge future research on ghost lineages and the potential for ancient introgression to drive some of the topological incongruence in Anthozoa. Notably, the deep divergences of sea anemone groups within hexacorals and the deep divergences within both orders of octocorals were the most unstable nodes and require further scrutiny.

The slow-evolutionary rate of mt genomes in anthozoans [24, 79] could also be partly responsible for the extreme mito-nuclear discordance seen here. Shearer et al. [79] hypothesized that background selection is influencing the slow substitution rates within mt genomes of anthozoans. Due to non-recombining mt loci, selection reduces variation not only at sites under selection, but at those that are linked as well [85]. We found that the mt genome is under strong purifying selection in both Hexacorallia and Octocorallia, with omega values close to zero in both classes. Another recent study found that some genes are under relaxed purifying selection in deep-sea taxa, with some sites in particular genes under positive selection [86]. We were not able to test for selection on nuclear loci, as none were in correct reading frames. However, because of the large number of loci used, we would not anticipate that all or even most nuclear loci would evolve under the same type of selection.

### Summary

Our results have demonstrated extreme mito-nuclear discordance in Anthozoa. Overall, non-recombining mt genomes that do not evolve neutrally and are likely to rapidly introgress are most likely influencing our ability to reconstruct accurate species relationships. Other studies have cautioned against the use of mtDNA for resolving phylogenetic relationships in anthozoans [21, 41] and even more broadly in metazoans [1], but unequal taxon sampling and non-matching tips have always been potential confounding issues in mito-nuclear comparisons. We included the same tips in the mt and nuclear phylogenies and sampled widely across all orders. Nonetheless, it is still possible that inadequate taxon sampling could influence the patterns of mito-nuclear discordance we observed, and that including more taxa in particular regions of the trees would stabilize some relationships. Even so, mito-nuclear discordance in hexacorals and octocorals is not an artifact of biased and incomparable taxon sampling, but instead, a signal of evolutionary processes that have shaped the genetic diversity of Anthozoa.

## Acknowledgements

This project was supported by the National Science Foundation (grants DEB-1457817 and DEB-1457581 to CSM and ER). P. Cowman provided the mt genome data for one specimen. A. Poliseno, L. Gusmão, and M. Xiao provided preliminary alignments of Zoantharia, Actiniaria, and Antipatharia. We thank D. Duchêne for providing advice on results of saturation tests.

## Author Contributions

AMQ, CSM, and ER conceived the study. AMQ conducted phylogenetic analyses, generated tables and figures, and along with CSM wrote the manuscript. AMQ, CSM, KS, RPR, IGLS, JH, and HHW assembled, annotated and aligned mitochondrial genes. NIR conducted selection tests. CSM, IGLS, NIR, ER, and HHW edited the manuscript. All authors approved the final version.

## Data Availability

Alignments, Tree files and code can be found on figshare XXXX

Mitochondrial genomes have been uploaded to genbank and respective numbers can be found in supplemental tables.

**Supplemental Figure 1.**
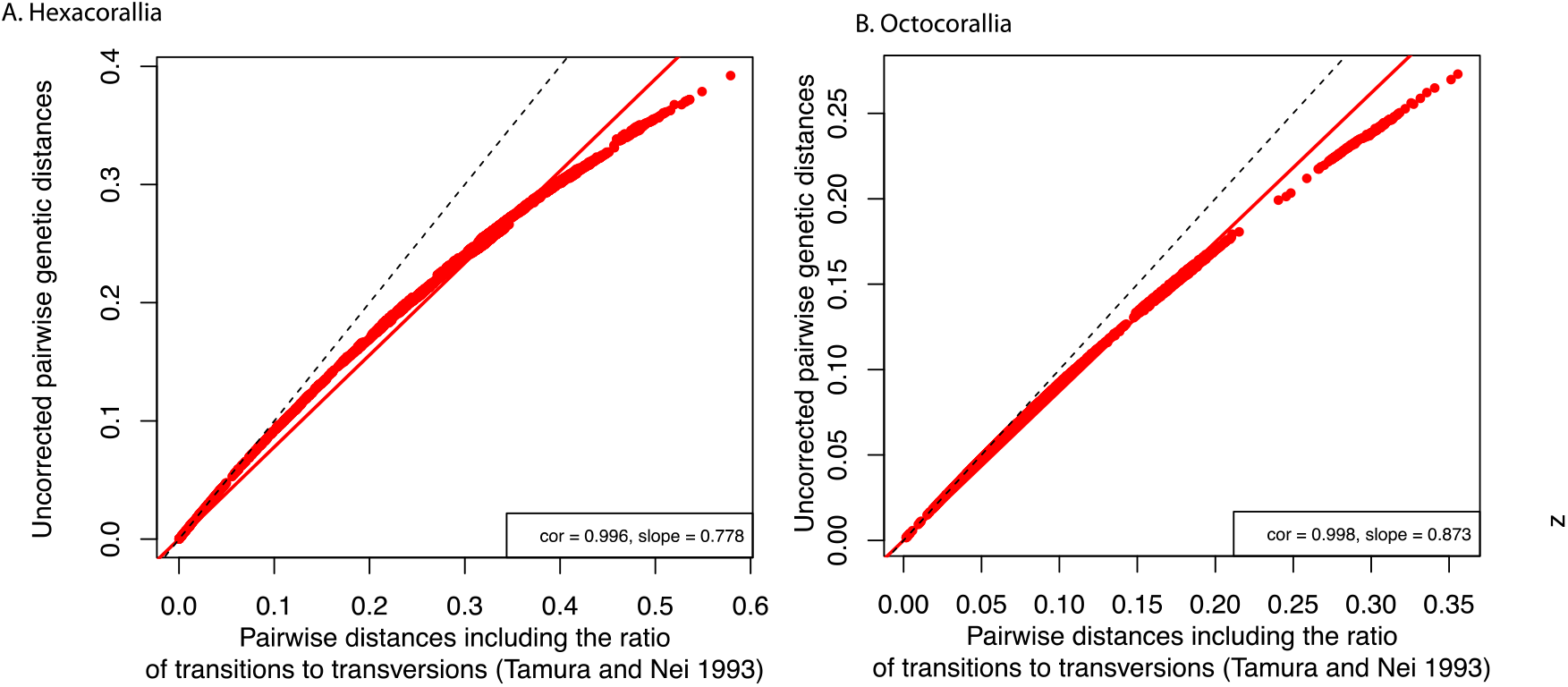
Saturation test results produced by PhyloMad for (A) Hexacorallia and (B) Octocorallia.

## References

1. Ballard, J. W. O., & Whitlock, M. C. The incomplete natural history of mitochondria. Mol. Ecol. 13(4), 729–744 (2004).

2. Janiak, M. C. et al. 205 newly assembled mitogenomes provide mixed evidence for rivers as drivers of speciation for Amazonian primates. Mol. Ecol. (2022)

3. Allio, R. et al. MitoFinder: efficient automated large-scale extraction of mitogenomic data in target enrichment phylogenomics. Mol. Ecol. Res., 20(4), 892–905 (2020)

4. Xiao, M et al. Mitogenomics suggests a sister relationship of Relicanthus daphneae (Cnidaria: Anthozoa: Hexacorallia: incerti ordinis) with Actiniaria. Sci. Rep., 9(1), 1–10. (2019).

5. Brown, W.M., George, M. & Wilson, A. C. Rapid evolution of animal mitochondrial DNA. Proc. Natl. Acad. Sci. U.S.A. 76, 1967–1971 (1979).

6. Platt, R. N. et al. Conflicting evolutionary histories of the mitochondrial and nuclear genomes in new world Myotis bats. Syst. Biol. 67, 236–249 (2018).

7. Tamashiro, R. A. et al. What are the roles of taxon sampling and model fit in tests of cyto-nuclear discordance using avian mitogenomic data? Mol. Phylogenet. Evol. 130, 132–142 (2019).

8. Kimball, R. T., Guido, M., Hosner, P. A. & Braun, E.L. When good mitochondria go bad: Cyto-nuclear discordance in landfowl (Aves: Galliformes). Gene. 801, 145841 (2021).

9. Lavrov, D. V. & Pett, W. Animal mitochondrial DNA as we do not know it: mt-genome organization and evolution in nonbilaterian lineages. Genome. Biol. Evol. 8, 2896–2913 (2016).

10. Beagley, C. T., Okada, N. A., & Wolstenholme, D. R. (1996). Two mitochondrial group I introns in a metazoan, the sea anemone Metridium senile: one intron contains genes for subunits 1 and 3 of NADH dehydrogenase. PNAS, 93(11), 5619–5623.

11. van Oppen, M. J., Catmull, J., McDonald, B. J., Hislop, N. R., Hagerman, P. J., & Miller, D. J. The mitochondrial genome of Acropora tenuis (Cnidaria; Scleractinia) contains a large group I intron and a candidate control region. J. Mol. Evol. 55(1), 1 (2002).

12. Medina, M., Collins, A. G., Takaoka, T. L., Kuehl, J. V. & Boore, J. L. Naked corals: Skeleton loss in Scleractinia. PNAS. 103, 9096–9100 (2006).

13. Fukami, H., Chen, C. A., Chiou, C. Y., & Knowlton, N. Novel group I introns encoding a putative homing endonuclease in the mitochondrial cox1 gene of Scleractinian corals. J. Mol. Evol. 64, 591–600 (2007).

14. Brugler, M. R., & France, S. C. The complete mitochondrial genome of the black coral Chrysopathes formosa (Cnidaria: Anthozoa: Antipatharia) supports classification of antipatharians within the subclass Hexacorallia. Mol. Phylogenet. Evol. 42, 776–788 (2007).

15. Stampar, S. N. et al. Linear mitochondrial genome in Anthozoa (Cnidaria): A case study in Ceriantharia. Sci Rep. 9, 6094 (2019).

16. Muthye, V., Mackereth, C. D., Stewart, J. B. & Lavrov, D. V. Large dataset of octocoral mitochondrial genomes provides new insights into mt-mutS evolution and function. DNA Repair (Amst). 110 (2022).

17. Bilewitch, J. P., & Degnan, S. M. A unique horizontal gene transfer event has provided the octocoral mitochondrial genome with an active mismatch repair gene that has potential for an unusual self-contained function. BMC Evol. Biol., 11, 1–15 (2011)

18. Hogan, R. I., Hopkins, K., Wheeler, A. J., Allcock, A. L. & Yesson, C. Novel diversity in mitochondrial genomes of deep-sea Pennatulacea (Cnidaria: Anthozoa: Octocorallia). Mitochondrial DNA Part A. 30, 764–777 (2019).

19. Uda, K. et al. Complete mitochondrial genomes of two Japanese precious corals, Paracorallium japonicum and Corallium konojoi (Cnidaria, Octocorallia, Coralliidae): Notable differences in gene arrangement. Gene. 476, 27–37 (2011).

20. Brockman, S. A. & McFadden, C. S. The mitochondrial genome of Paraminabea aldersladei (Cnidaria: Anthozoa: Octocorallia) supports intramolecular recombination as the primary mechanism of gene rearrangement in octocoral mitochondrial genomes. Genome. Biol. Evol. 4, 994–1006 (2012).

21. Figueroa, D. F. & Baco, A. R. Octocoral mitochondrial genomes provide insights into the phylogenetic history of gene order rearrangements, order reversals, and cnidarian phylogenetics. Genome. Biol. Evol. 7, 391–409 (2015).

22. Brown, W. M., Prager, E. M., Wang, A. & Wilson, A. C. Mitochondrial DNA sequences of primates: Tempo and mode of evolution. J. Mol. Evol. 18, 225–239 (1982).

23. Vawter, L. & Brown, W. M. Nuclear and mitochondrial DNA comparisons reveal extreme rate variation in the molecular clock. Science. 234, 194–196 (1986).

24. Hellberg, M. E. No variation and low synonymous substitution rates in coral mtDNA despite high nuclear variation. BMC Evol. Biol. 6, 24 (2006).

25. Shearer, T. L. & Coffroth, M. A. DNA Barcoding: Barcoding corals: limited by interspecific divergence, not intraspecific variation. Mol. Ecol Res. 8, 247–255 (2008).

26. Huang, D., Meier, R., Todd, P. A. & Chou, L. M. Slow mitochondrial COI sequence evolution at the base of the metazoan tree and its implications for DNA barcoding. J. Mol. Evol. 66, 167–174 (2008).

27. Quattrini, A. M. et al. Palaeoclimate ocean conditions shaped the evolution of corals and their skeletons through deep time. Nat. Ecol. Evol. 4, 1531–1538 (2020).

28. McFadden, C. S. et al. Phylogenomics, origin, and diversification of anthozoans (Phylum Cnidaria). Syst Biol. 70, 635–647 (2021).

29. Park, E. et al. Estimation of divergence times in cnidarian evolution based on mitochondrial protein-coding genes and the fossil record. Mol. Phylogenet. Evol. 62, 329–345 (2012).

30. Kayal, E., Roure, B., Philippe, H., Collins, A. G. & Lavrov, D. V. Cnidarian phylogenetic relationships as revealed by mitogenomics. BMC Evol. Biol. 13, 5 (2013).

31. Daly, M. et al. The phylum Cnidaria: A review of phylogenetic patterns and diversity 300 years after Linnaeus. Zootaxa. 1668, 127–182 (2007).

32. Zapata, F. et al. Phylogenomic analyses support traditional relationships within Cnidaria. PLoS One. 10, e0139068. 10.1371/journal.pone.0139068. (2015).

33. Kayal, E. et al. Phylogenomics provides a robust topology of the major cnidarian lineages and insights on the origins of key organismal traits. BMC Evol. Biol. 18, 68 (2018).

34. Osigus, H-J., Eitel, M., Bernt, M., Donath, A. & Schierwater, B. Mitogenomics at the base of metazoa. Mol. Phylogenet. Evol. 69, 339–351 (2013).

35. Stampar, S. N., Maronna, M. M., Kitahara, M. V., Reimer, J. D. & Morandini, A. C. Fast-evolving mitochondrial DNA in Ceriantharia: A reflection of Hexacorallia paraphyly? PLoS One. 9, e86612. 10.1371/journal.pone.0086612 (2014).

36. Kitahara, M. V. et al. The “Naked Coral” hypothesis revisited – Evidence for and against Scleractinian monophyly. PLoS One. 9, e94774. 10.1371/journal.pone.0094774 (2014).

37. Seiblitz, I. G. L. et al. The earliest diverging extant scleractinian corals recovered by mitochondrial genomes. Sci. Rep. 10, 20714; 10.1038/s41598-020-77763-y (2020).

38. Stolarski, J. et al. A. A modern scleractinian coral with a two-component calcite– aragonite skeleton. PNAS, 118(3), e2013316117 (2021).

39. McFadden, C. S., France, S. C., Sánchez, J. A. & Alderslade, P. A molecular phylogenetic analysis of the Octocorallia (Cnidaria: Anthozoa) based on mitochondrial protein-coding sequences. Mol. Phylogenet. Evol. 41, 513–527 (2006).

40. McFadden, C. S., van Ofwegen, L. P. & Quattrini, A. M. Revisionary systematics of Octocorallia (Cnidaria: Anthozoa) guided by phylogenomics. Bull. Syst. Biol. 1, 8735; https://doi.org/10.18061/bssb.v1i3.8735 (2022).

41. Pratlong, M., Rancurel, C., Pontarotti, P. & Aurelle, D. Monophyly of Anthozoa (Cnidaria): Why do nuclear and mitochondrial phylogenies disagree? Zool Scr. 46, 363–371 (2016).

42. Quattrini, A. M., et al. Universal target-enrichment baits for anthozoan (Cnidaria) phylogenomics: New approaches to long-standing problems. Mol. Ecol. Resour. 18, 281–295 (2018).

43. Faircloth, B. C. Illumiprocessor: a trimmomatic wrapper for parallel adapter and quality trimming. 10.6079/J9ILL (2013).

44. Bolger, A. M., Lohse, M. & Usadel, B. Trimmomatic: a flexible trimmer for Illumina sequence data. Bioinformatics. 30, 2114–2120 (2014).

45. Bankevich, A. et al. SPAdes: A new genome assembly algorithm and its applications to single-cell sequencing. J. Comput. Biol. 19, 455–477 (2012).

46. Haas, B. J. et al. De novo transcript sequence reconstruction from RNA-seq using the Trinity platform for reference generation and analysis. Nat Protoc. 8, 1494–1512 (2013).

47. Katoh, K. & Standley, D. M. MAFFT multiple sequence alignment software version 7: Improvements in performance and usability. Mol Biol Evol. 30, 772–780 (2013).

48. Dierckxsens, N., Mardulyn, P. & Smits, G. NOVOPlasty: de novo assembly of organelle genomes from whole genome data. Nucleic Acids Res. 45, e18. 10.1093/nar/gkw955 (2016).

49. Donath, A. et al. Improved annotation of protein-coding genes boundaries in metazoan mitochondrial genomes. Nucleic Acids Res. 47, 10543–10552 (2019).

50. Duchêne, D. A., Mather, N., van der Wal, C. & Ho, S. Y. W. Excluding loci with substitution saturation improves inferences from phylogenomic data. Syst Biol. 71, 676–689 (2021).

51. Duchêne, D. A., Duchêne, S. & Ho, S. Y. W. PhyloMAd: efficient assessment of phylogenomic model adequacy. Bioinformatics. 34, 2300–2301 (2018).

52. Yang, Z. PAML 4: Phylogenetic analysis by maximum likelihood. Mol. Biol. Evol. 24, 1586–1591 (2007).

53. Nguyen, L-T., Schmidt, H. A., von Haeseler, A. & Minh, B. Q. IQ-TREE: A fast and effective stochastic algorithm for estimating maximum-likelihood phylogenies. Mol. Biol. Evol. 32, 268–274 (2015).

54. Kalyaanamoorthy, S., Minh, B. Q., Wong, T. K. F., von Haeseler, A. & Jermiin, L. S. ModelFinder: fast model selection for accurate phylogenetic estimates. Nat Methods. 14, 587–589 (2017).

55. Hoang, D. T., Chernomor, O., von Haeseler, A., Minh, B. Q. & Vinh, L. S. UFBoot2: Improving the ultrafast bootstrap approximation. Mol. Biol. Evol. 35, 518–522 (2018).

56. Anisimova, M., Gil, M., Dufayard, J-F., Dessimoz, C. & Gascuel, O. Survey of branch support methods demonstrates accuracy, power, and robustness of fast likelihood-based approximation schemes. Syst. Biol. 60, 685–699 (2011).

57. Minh, B. Q., Hahn, M. W. & Lanfear, R. New methods to calculate concordance factors for phylogenomic datasets. Mol. Biol. Evol. 37, 2727–2733 (2020).

58. Zhang, C., Rabiee, M., Sayyari, E. & Mirarab, S. ASTRAL-III: polynomial time species tree reconstruction from partially resolved gene trees. BMC Bioinformatics. 19, 153 (2018).

59. Revell, L. J. phytools: an R package for phylogenetic comparative biology (and other things). Methods. Ecol. Evol. 2, 217–223 (2012).

60. Mai, U. & Mirarab, S. TreeShrink: fast and accurate detection of outlier long branches in collections of phylogenetic trees. BMC Genomics. 19, 272 (2018).

61. Robinson, D. F. & Foulds, L. R. Comparison of phylogenetic trees. Math Biosci. 53, 131–147 (1981).

62. Poliseno, A. et al. Evolutionary implications of analyses of complete mitochondrial genomes across order Zoantharia (Cnidaria: Hexacorallia). J. Zool. Syst. Evol. Res. 58, 858–868 (2020).

63. Emblem, Å. et al. Sea anemones possess dynamic mitogenome structures. Mol. Phylogenet. Evol. 75, 184–193 (2014).

64. Celis, J. S. et al. Evolutionary and biogeographical implications of degraded LAGLIDADG endonuclease functionality and group I intron occurrence in stony corals (Scleractinia) and mushroom corals (Corallimorpharia). PLoS One, 12, e0173734 (2017).

65. Foox, J., Brugler, M., Siddall, M. E., & Rodríguez, E. Multiplexed pyrosequencing of nine sea anemone (Cnidaria: Anthozoa: Hexacorallia: Actiniaria) mitochondrial genomes. Mitochondrial DNA A 27, 2826–2832 (2016).

66. Seiblitz, I. G. et al. Caryophylliids (Anthozoa, Scleractinia) and mitochondrial gene order: Insights from mitochondrial and nuclear phylogenomics. Mol. Phyl. Evol. 175, 107565 (2022).

67. Brugler, M. R., & France, S. C. The mitochondrial genome of a deep-sea bamboo coral (Cnidaria, Anthozoa, Octocorallia, Isididae): genome structure and putative origins of replication are not conserved among octocorals. J. Mol. Evol, 67(2), 125–136.

68. Johansen, S. D., & Emblem, Å. Mitochondrial Group I introns in hexacorals are regulatory genetic elements In Advances in the Studies of the Benthic Zone (IntechOpen) 101 (2020).

69. DeBiasse, M., et al. A cnidarian phylogenomic tree fitted with hundreds of 18S leaves. bioRxiv. (2022)

70. Rodríguez, E., et al. Hidden among sea anemones: the first comprehensive phylogenetic reconstruction of the order Actiniaria (Cnidaria, Anthozoa, Hexacorallia) reveals a novel group of hexacorals. PloS one, 9(5), e96998 (2014).

71. van Oppen, M. V., Willis, B. L., Vugt, H. V., & Miller, D. J. Examination of species boundaries in the Acropora cervicornis group (Scleractinia, Cnidaria) using nuclear DNA sequence analyses. Mol. Ecol. 9(9), 1363–1373 (2000).

72. Vollmer, S. V., & Palumbi, S. R. Hybridization and the evolution of reef coral diversity. Science, 296(5575), 2023–2025. (2002)

73. Reimer, J. D., Takishita, K., Ono, S., Tsukahara, J., & Maruyama, T. Molecular evidence suggesting interspecific hybridization In Zoanthus spp. (Anthozoa: Hexacorallia). Zool. Sci. 24, 346–359 (2007).

74. Combosch, D. J., & Vollmer, S. V. Trans-Pacific RAD-Seq population genomics confirms introgressive hybridization in Eastern Pacific Pocillopora corals. Mol. Phylogenet. Evol. 88, 154–162 (2015).

75. Quattrini, A. M. et al. A next generation approach to species delimitation reveals the role of hybridization in a cryptic species complex of corals. BMC Evol. Biol. 19, 1–19 (2019).

76. Hellberg, M. E., Prada, C., Tan, M. H., Forsman, Z. H., & Baums, I. B. Getting a grip at the edge: recolonization and introgression in eastern Pacific Porites corals. J. Biogeogr. 43, 2147–2159 (2016).

77. Forsman, Z. H. et al. Coral hybridization or phenotypic variation? Genomic data reveal gene flow between Porites lobata and P. compressa. Mol. Phylogenet. Evol. 111, 132–148 (2017).

78. Chan, K. M., & Levin, S. A. Leaky prezygotic isolation and porous genomes: rapid introgression of maternally inherited DNA. Evol. 59, 720–729 (2005).

79. Shearer, T. L., Van Oppen, M. J. H., Romano, S. L., & Wörheide, G. Slow mitochondrial DNA sequence evolution in the Anthozoa (Cnidaria). Mol. Ecol. 11, 2475–2487 (2002).

80. Hey, J. Isolation with migration models for more than two populations. Mol. Biol. Evol. 27(4), 905–920. (2010)

81. Gompert, Z., Forister, M. L., Fordyce, J. A., & Nice, C. C. Widespread mito-nuclear discordance with evidence for introgressive hybridization and selective sweeps In Lycaeides. Mol. Ecol. 17(24), 5231–5244 (2008).

82. Linnen, C. R., & Farrell, B. D. Mitonuclear discordance is caused by rampant mitochondrial introgression In Neodiprion (Hymenoptera: Diprionidae) sawflies. Evolution: International Journal of Organic Evolution, 61(6), 1417–1438 (2007).

83. Hibbins, M. S., & Hahn, M. W. Phylogenomic approaches to detecting and characterizing introgression. Genetics 220, iyab173 (2022).

84. Li, Y., & Wu, D. D. Finding unknown species in the genomes of extant species. J. Genetics Genomics. 48(10), 867–871 (2021)

85. Schrider, D. R. Background selection does not mimic the patterns of genetic diversity produced by selective sweeps. Genetics 216, 499–519 (2020).

86. Ramos, N.I., DeLeo D.M., McFadden, C.S., & Quattrini, A.M. Selection in coral mitogenomes, with insights into adaptations in the deep sea. Sci. Rep. Submitted.

