## Supplementary figures and images for "Extreme mito-nuclear discordance within Anthozoa, with notes on unique properties of their mitochondrial genomes"

### Supplemental Figure 1

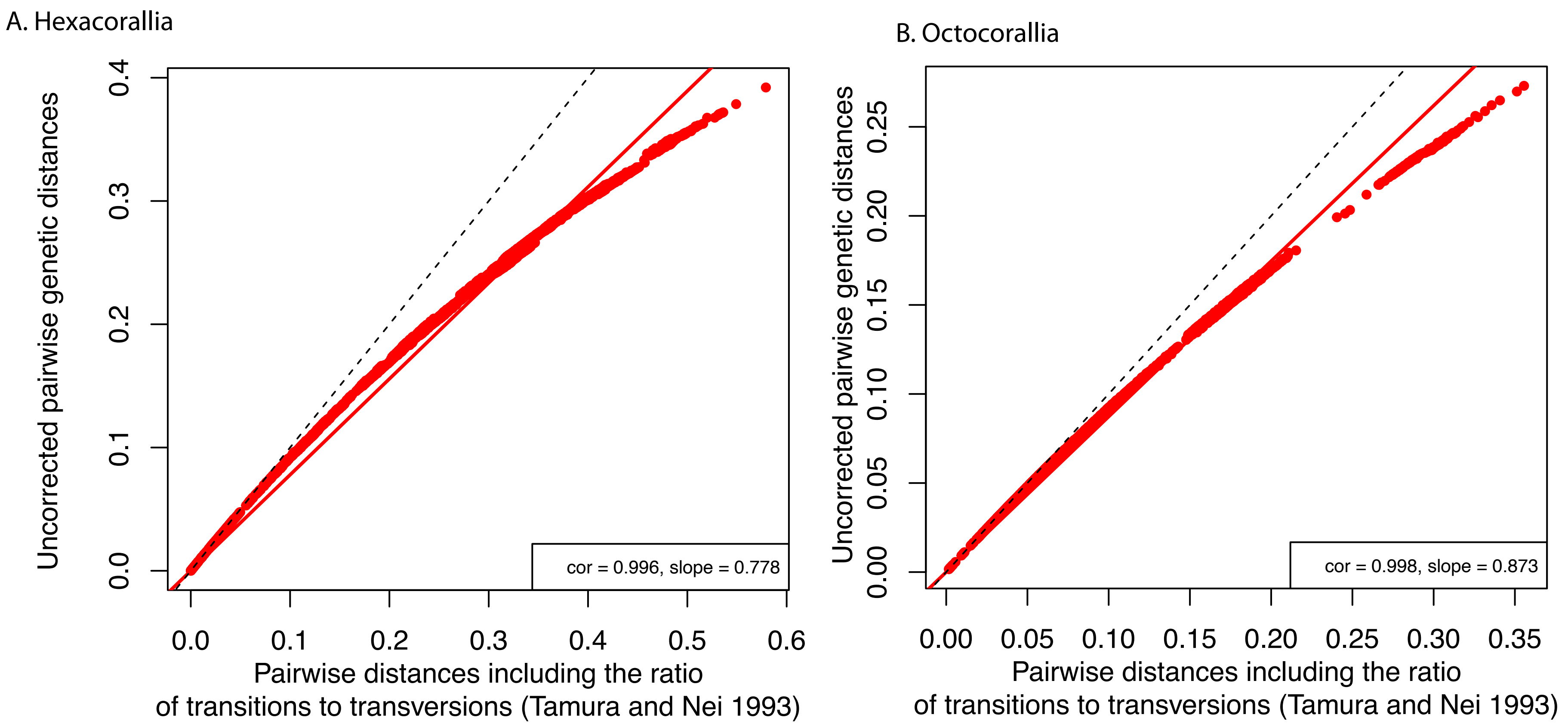
